# Low circulating choline, a modifiable dietary factor, is associated with the pathological progression and metabolome dysfunction in Alzheimer’s disease

**DOI:** 10.1101/2023.05.06.539713

**Authors:** Jessica M. Judd, Paniz Jasbi, Wendy Winslow, Geidy E. Serrano, Thomas G. Beach, Judith Klein-Seetharaman, Ramon Velazquez

## Abstract

Most Americans (∼90%) are deficient in dietary choline, an essential nutrient. Associations between circulating choline and pathological progression in Alzheimer’s disease (AD) remain unknown. Here, we examined these associations and performed a metabolomic analysis in blood serum from severe AD, moderate AD, and healthy controls. Additionally, to gain mechanistic insight, we assessed the effects of dietary choline deficiency (Ch-) in 3xTg-AD mice and choline supplementation (Ch+) in APP/PS1 mice. In humans, we found AD-associated reductions in choline, it’s derivative acetylcholine (ACh), and elevated pro-inflammatory cytokine TNFα. Choline and ACh were negatively correlated with Plaque density, Braak stage, and TNFα, but positively correlated with MMSE and brain weight. Metabolites L-Valine, 4-Hydroxyphenylpyruvic, Methylmalonic, and Ferulic acids were associated with choline levels. In mice, Ch-paralleled AD severe, but Ch+ was protective. In conclusion, low circulating choline is associated with AD-neuropathological progression, illustrating the importance of dietary choline consumption to offset disease.

## Introduction

Alzheimer’s disease (AD) is a neurodegenerative disorder that currently affects 6.5 million people aged 65 and older, with an estimated increase to 12.7 million by 2050^1^. AD is characterized by the presence of amyloid beta (Aβ) plaques and neurofibrillary tau tangles (NFT) that result in progressive loss of memory and other cognitive abilities^2^. While clinical symptomologies of AD typically appear later in life, there is a long preclinical phase, where molecular dysregulation, including elevations in neuroinflammatory factors, such as tumor necrosis factor alpha (TNFα), contribute to pathogenesis^3^. A better understanding of preclinical processes could prevent or delay disease development.

Environmental factors, including diet, may contribute to AD pathogenesis. Choline, a B-like micronutrient, plays key roles across a wide range of organ systems, including being a precursor of both acetylcholine (ACh) and cell-membrane lipids^4^. Only 30% of the required choline is produced endogenously by phosphatidylethanolamine-N-methyltransferase (PEMT) in the liver; the rest must be consumed^5^. Dietary guidelines by the Institute of Medicine are focused on liver health and recommend 550 and 425 mg/day for men and women, respectively, and 550mg/day for pregnant women given the fetus’s developmental requirements^6^. Normal circulating choline concentrations in fasted, non-pregnant humans is ∼9.56±0.49 μmol/L^4^. Circulating choline levels are controlled by choline supply and the ability of tissues to accumulate choline^7^. The rate of choline transport across the blood brain barrier is positively dependent on circulating choline concentration and requires choline transporters^7^. Dietary choline deficiency produces detrimental outcomes, including nonalcoholic fatty liver disease, glucose metabolism impairments, cardiovascular disease, and impaired cognition^5, 8^. Alarmingly, choline deficiency is observed worldwide; including ∼90% of Americans, with average dietary intake of males being 402mg/day, while females only consume 278mg/day^6, 9^. This is notable as choline modulates metabolic functions whose dysregulation increase AD prevalence^10^, and AD incidence is higher in females^2^.

Recent work highlights choline’s importance for healthy cognitive aging and that deficiency can contribute to AD development^11, 12^. Older adults who consume 187.6–399.5 mg/day of choline have less cognitive decline than those consuming <187.6 mg/day^11^. Additionally, low dietary choline intake across the lifespan increases the risk of dementia (< 215mg/day)^12^. However, these reports are based on dietary supplementation questionnaire estimates, and not on measured circulating choline. Further, abnormal PEMT increases AD incidence^13^ and reduction in ACh is a well-documented disease feature of AD^14^. AD mouse models support the importance of dietary choline for healthy aging; dietary choline above the recommended daily amount protects against cognitive decline and AD pathology^15^, while choline deficiency exacerbates pathology^8^. Moreover, metabolomic studies show that metabolic changes accompany neurological disorders ^16, 17^ and have identified differences in CSF choline levels between healthy controls, and a group of mild cognitive impairment (MCI) and AD patients^16^. However, metabolite dysregulation in AD related to reductions in choline has not been elucidated and may reveal novel disease mechanisms and biomarkers of disease progression.

The goal of the current study is to gain insight into the circulatory levels of choline as a function of disease progression, using blood serum from moderate and severe stage AD patients and healthy controls to understand its association with AD symptomology. Additionally, we examined the role of dietary choline deficiency and supplementation in two mouse models of AD to parallel conditions observed in human AD patients. We also identified metabolites in human serum that are associated with choline levels. We hypothesize that reduced circulating choline levels will be associated with increased AD pathologies, while supplementation with choline will reduce neuropathology.

## Results

### Pathological AD hallmarks in human subjects

Human serum samples were obtained from the Arizona Study of Aging and Neurodegenerative Disorders/Brain and Body Donation Program^18, 19^. AD Moderate (AD Mod) and AD Severe (AD Sev) corresponded with NIA-RI Intermediate and High classifications, respectively^19, 20^. Healthy controls (CON) did not meet the criteria for AD. We assessed human subjects on profile metrics (Supplementary Table 1). Analysis between the AD Mod and AD Sev showed no significant differences for years since diagnosis (Supplemental Figure 1A; *t*_22_ = 1.046, *P* = 0.3068) or age at diagnosis (Supplemental Fig. 1B; *t*_22_ = 1.300, *P* = 0.2069). There were no significant differences between the groups (CON, AD Mod, AD Sev; Supplemental Fig. 1C-F) for final BMI (*F*_2,28_ = 0.5525, *P* = 0.5817), time since last BMI (*F*_2,28_ = 0.5406, *P* = 0.5884), expired age (*F*_2,33_ = 0.2349, *P* = 0.7920), and PMI tissue collection (*F*_2, 33_ = 0.8296, *P* = 0.4451).

We next analyzed the three groups (CON, AD Mod, AD Sev) to confirm differences in AD progression. There were significant group differences in cognition, as measured by the Mini Mental State Exam (MMSE; Supplemental Fig. 1G; *F*_2,26_ = 15.68, *P* < 0.0001). AD Sev exhibited lower scores than CON (*P* < 0.0001) and AD Mod (*P* = 0.0059). For CERAD neuritic plaque density, a semi-quantitative measure of deteriorating neuronal material surrounding Aβ plaques, we found significant group differences (Supplemental Fig. 1H; *F*_2,33_ = 113.3, *P* < 0.0001). CON had significantly less plaque density than AD Mod (*P* < 0.0001) and AD Sev (*P* < 0.0001). AD Mod trended towards less plaque density than AD Sev (*P* = 0.0625). For Braak score, a measure of the extent of neurofibrillary tau tangle pathology, we found significant group differences (Supplemental Fig. 1I; *F*_2,33_ = 200.2, *P* < 0.0001). AD Sev had higher Braak stage classification than both CON (P < 0.0001) and AD Mod (P < 0.0001); AD Mod had higher Braak stage classification than CON (*P* < 0.0001). Lastly, we analyzed brain weight and found significant group differences (Supplemental Fig. 1J; *F*_2,28_ = 7.825, *P* = 0.0016). AD Sev brains weighed significantly less than CON (*P* = 0.0012). Collectively, these results highlight disease associated changes between the AD Mod and AD Sev compared to CON subjects.

### Serum choline, acetylcholine, and TNFα levels differ between control and AD groups, correlating with neuropathology

We assessed serum levels of choline, ACh, and TNFα. We found significant group differences of serum choline levels (Fig. 1A; *F*_2,33_ = 62.45, *P* < 0.0001); both AD Mod (*P* < 0.0001) and AD Sev (*P* < 0.0001) had lower choline than CON. AD Sev had lower choline than AD Mod (*P* = 0.0155). For serum ACh, we found significant group differences (Fig. 1B; *F*_2,33_ = 204.2, *P* < 0.0001); both AD Mod (*P* < 0.0001) and AD Sev (*P* < 0.0001) had lower levels than CON. Additionally, AD Sev had lower ACh than AD Mod (*P* < 0.0001). For TNFα, we found significant group differences (Fig. 1C; *F*_2,33_ = 18.82, *P* < 0.0001); AD Mod (*P* = 0.0231) and AD Sev (*P* < 0.0001) had higher levels than CON. Additionally, AD Sev had higher serum TNFα than AD Mod (*P* = 0.0071). These results highlight a disease-associated reduction of choline and ACh, and an increase in TNFα levels in AD patients.

Next, we performed correlations between serum choline, ACh, TNFα, MMSE scores, CERAD neuritic plaque density, Braak stage, and brain weight. We found significant positive correlations between choline and ACh (Fig. 1D; *r*_34_ = 0.7713, *P* < 0.001), and between both choline (Fig. 1E; *r*_27_ = 0.5936, *P* = 0.0007), ACh (Fig. 1F; *r*_27_ = 0.7064, *P* < 0.0001) and MMSE, indicating that higher choline and ACh are associated with higher MMSE outcomes. Conversely, TNFα was negatively correlated with MMSE (Fig. 1G; *r*_27_ = -0.5275, *P* = 0.0033). We next performed correlations between CERAD neuritic plaque density and choline, ACh, and TNFα. We found significant negative correlations between both choline (Fig. 1H; *r*_34_ = -0.6674, *P* < 0.0001), ACh (Fig. 1I; *r*_34_ = -0.8884, *P* < 0.0001) and plaque density; less choline and ACh was associated with higher plaque load. Further, a positive correlation showed that as TNFα increased, so did plaque density (Fig. 1J; *r*_34_ = 0.5858, *P* = 0.0002).

**Figure 1:**
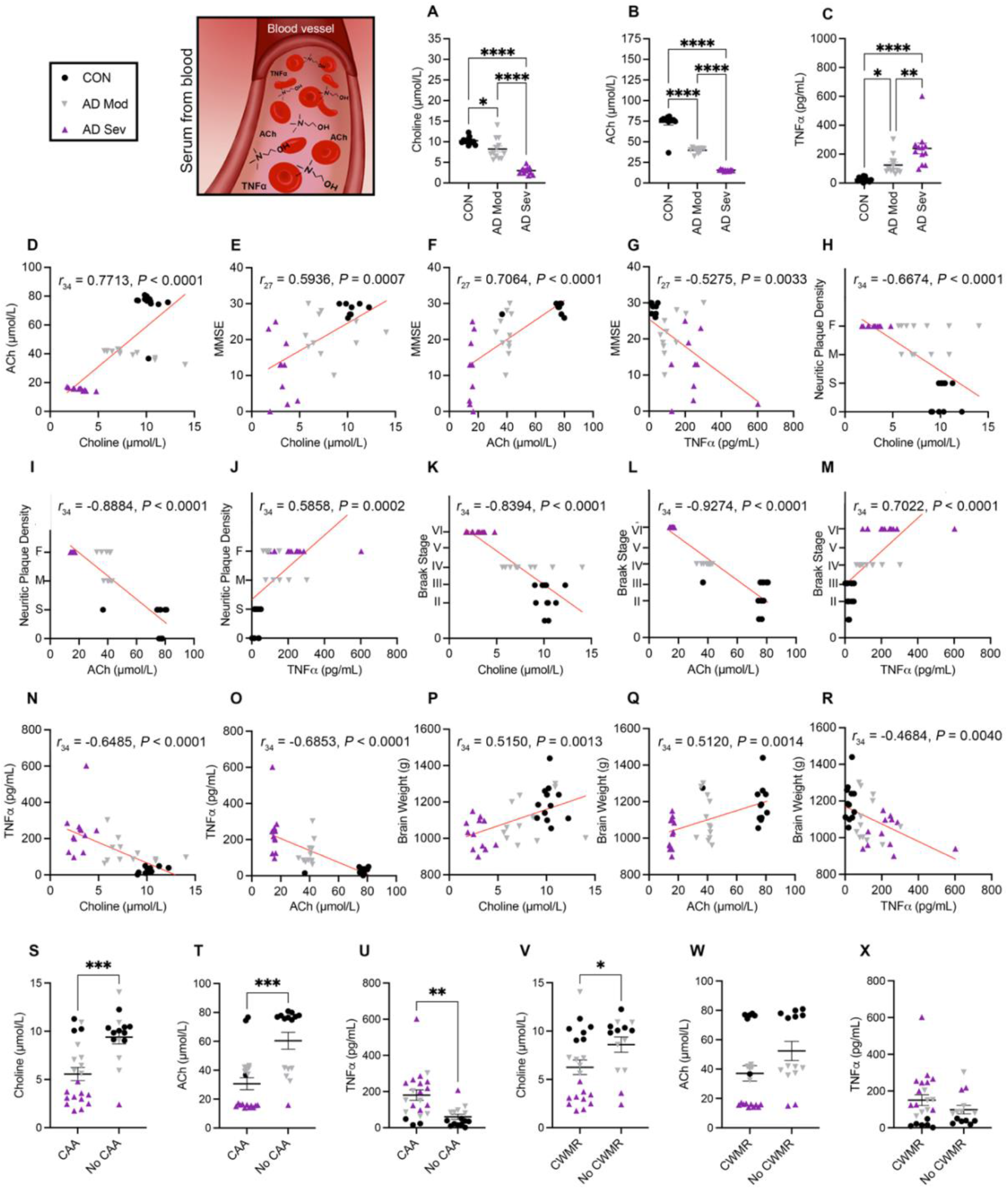
Serum free choline, ACh and TNFα levels correlate with AD-associated pathologies. **(A-C)** Serum levels of free choline, ACh, and TNFα. **(D)** Correlation between choline and ACh. **(E-G)** Correlations with MMSE and serum measurements. **(H-J)** Correlations with CERAD neuritic plaque density and serum measurements. 0=Zero, S=Sparce, M=Moderate, F=Frequent. **(K-M)** Correlations between Braak stage and serum measurements. **(N-O)** Correlation between choline, ACh and TNFα. **(P-R)** Correlations between brain weight and serum measurements. Choline, ACh, and TNFα levels differ in cases with cerebral amyloid angiopathy (CAA) and cerebral white matter rarefaction (CWMR). **(S-U)** Choline and ACh are significantly reduced in CAA positive cases, while TNFα is increased. **(V-X)** Choline is significantly reduced in cases with CWMR, but ACh and TNFα are not significantly different between CWMR cases. Data are reported as means ± SEM. *P < 0.05, **P < 0.01, ***P < 0.001.

Next, we performed correlations between serum choline, ACh, TNFα, and Braak stage. We found highly significant negative correlations between choline (Fig. 1K; *r*_34_ = -0.8394, *P* < 0.0001), ACh (Fig. 1L; *r*_34_ = -0.9274, *P* < 0.0001) and Braak stage. A significant positive correlation illustrated that as TNFα increase, so does Braak stage (Fig. 1M; *r*_34_ = 0.7022, *P* < 0.0001). Further, there were significant negative correlations between both choline (Fig. 1N; *r*_34_ = -0.6485, *P* < 0.0001), ACh (Fig. 1O; *r*_34_ = -0.6853, *P* < 0.0001) and TNFα. Lastly, there were significant positive correlations for both choline (Fig. 1P; *r*_34_ = 0.5150, *P* = 0.0013), ACh (Fig. 1Q; *r*_34_ = 0.5120, *P* = 0.0014) and brain weight, and a negative correlation between TNFα and brain weight (Fig. 1R; *r*_34_ = -0.4684, *P* = 0.0040). These results highlight that higher choline and ACh levels correlate with lower AD pathology, and elevated TNFα is associated with increased AD pathology.

### Choline, acetylcholine, and TNFα differ in cases with cerebral amyloid angiopathy (CAA) and cerebral white matter rarefaction (CWMR)

CAA is a condition where amyloid builds up in the walls of arteries and may lead to stroke and dementia^21^. To determine if choline, ACh and TNFα are significantly different in patients with confirmed CAA, we stratified human cases into those with and without CAA. We found that both choline (*t_34_* = 3.830*, P* = 0.0005) and ACh (*t_34_* = 4.257*, P* = 0.0002) are significantly lower in patients with CAA (Fig. 1S, T). We also found a significant increase of TNFα in patients with CAA (Fig. 1U; *t_34_* = 3.335, *P* = 0.0021). Given that choline is important in the biosynthesis of phosphatidylcholine, a major metabolite of myelin, we next stratified human cases into those with or without CWMR, which is a measure of overall white matter integrity and myelin loss^22^. We then analyzed choline, ACh and TNFα serum between the CWMR categorized groups. Interestingly, we found that patients with CWMR had lower choline (Fig. 1V; *t_34_* = 2.055, *P* = 0.0476), however, there were no significant differences in ACh or TNFα between patients with or without CWMR (Fig. 1W, X), suggesting that low circulating choline may be associated with a reduction in myelin, but not ACh nor TNFα.

### A choline deficient diet in mice induces a phenotype that parallels human AD

To determine the consequences of dietary choline deficiency, we placed 3xTg-AD and NonTg mice on either a choline normal (ChN) or choline deficient (Ch-) diet from three to 12 months of age (Fig. 2A). Blood plasma was obtained from live mice at 7 and 12 months of age to measure choline, ACh, and TNFα. We found a significant main effect of age; plasma choline was decreased with age (Fig. 2B; *F*_1,20_ = 158.3, *P* < 0.0001). At 7 months of age, there was a main effect of genotype (*F*_1,20_ = 198.1, *P* < 0.0001), with lower choline in the 3xTg-AD mice compared to the NonTg mice. Additionally, there was a significant main effect of diet (*F*_1,20_ = 40.75, *P* < 0.0001), with lower choline in the Ch-mice compared to the ChN counterparts. At 12 months, we found a significant main effect of diet, where 3xTg-AD Ch- and NonTg Ch-mice showed reduced choline in both plasma (*F*_1,20_ = 96.08, *P* < 0.0001) and cortex (Fig. 2C; *F*_1,20_ = 165.3, *P* < 0.0001). The 12-month choline findings were previously reported^8^ and are reiterated for clarity. For plasma ACh, we also found a significant main effect of age (Fig. 2D; *F*_1,20_ = 1335, *P* < 0.0001), where levels were decreased with age. At 7 months, we found significant main effects of both genotype (*F*_1,20_ = 348.2, *P* < 0.0001) and diet (*F*_1,20_ = 124.5, *P* < 0.0001), with lower ACh in 3xTg-AD mice and Ch-mice. Additionally, we found a significant genotype by diet interaction (*F*_1,20_ = 92.79, *P* < 0.0001); there was significantly less ACh in the 3xTg-AD Ch-mice compared to their ChN counterparts (*P* < 0.0001). There were no significant dietary differences in the NonTg mice. At 12 months, we found significant main effects for both genotype (*F*_1,20_ = 57.34, *P* < 0.0001) and diet (*F*_1,20_ = 22.77, *P* = 0.0001), where 3xTg-AD mice had lower ACh than NonTg and the Ch-diet led to reduced levels. We also found a significant genotype by diet interaction (*F*_1,20_ = 18.40, *P* = 0.0004), where NonTg Ch-mice had lower ACh than their ChN counterparts (*P* < 0.0001). Interestingly, both the 3xTg-AD ChN (*P* < 0.0001) and 3xTg-AD Ch-(*P* < 0.0001) mice had lower ACh than the NonTg ChN, indicating diet-independent reductions in 3xTg-AD mice. Taken together, a Ch-diet reduces plasma and cortical choline in both NonTg and 3xTg-AD mice. Interestingly, while only 3xTg-AD mice show substantial reduction of ACh following a Ch-at 7 months, by 12 months both 3xTg-AD mice groups and the NonTg Ch-mice show significantly reduced ACh.

**Figure 2:**
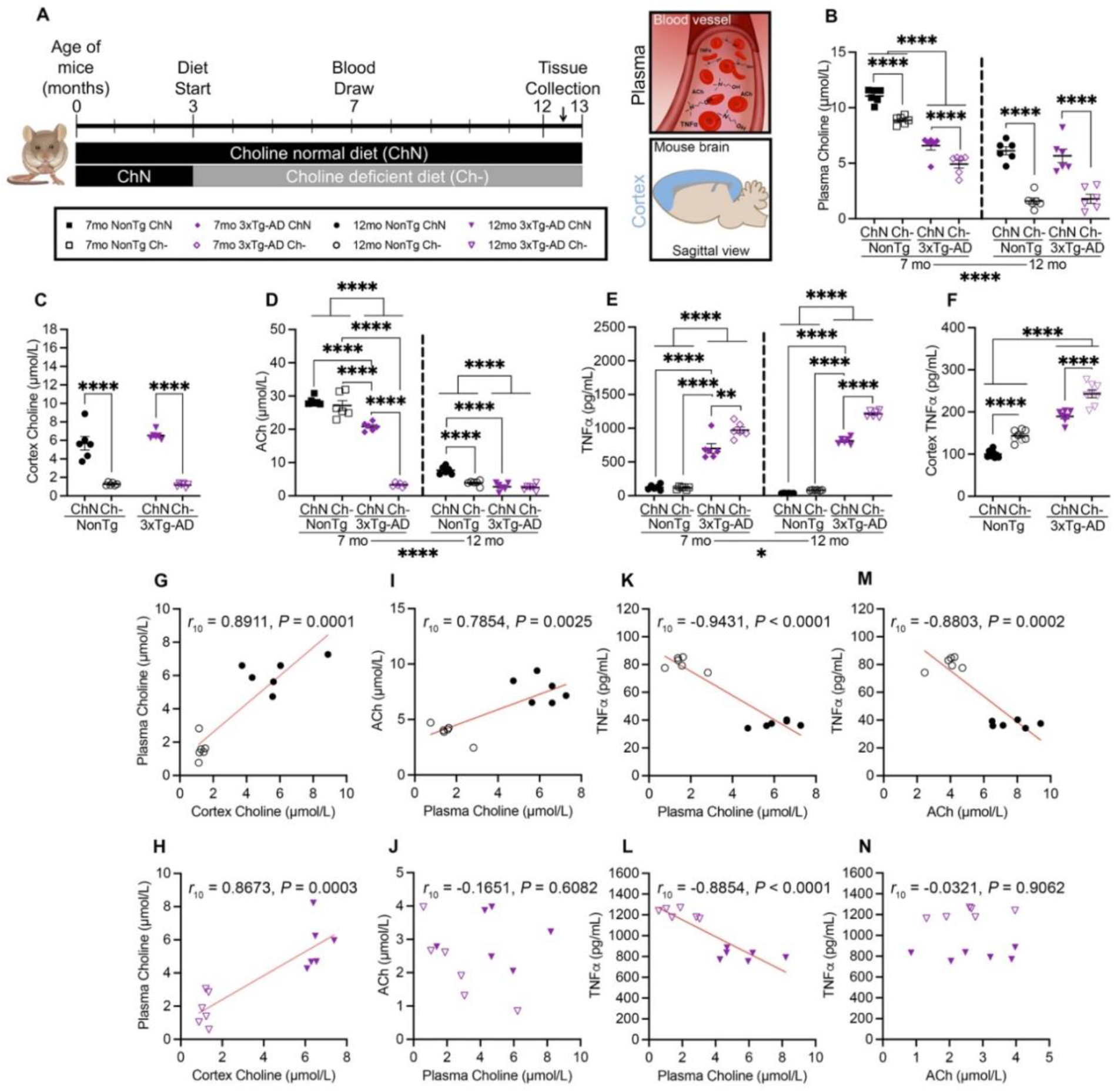
A choline deficiency (Ch-) diet results in reduced free choline and ACh, but elevated TNFα at 7 and 12 months (mo), Aβ pathological markers, and tau phosphorylation. **(A)** Timeline of dietary choline manipulation. (B) Ch-led to decreases in plasma choline at both 7 and 12 mo. At 7mo 3xTg-AD mice have lower plasma choline than NonTg mice. **(C)** At 12 mo, Ch-mice exhibited lower cortical choline levels. **(D)** At 7 mo, 3xTg-AD mice had lower plasma ACh than Non-Tg and the 3xTg-AD Ch-mice had the lowest plasma ACh. At 12 mo, the Ch-diet led to a decrease in ACh in NonTg mice and 3xTg-AD mice had lower ACh than NonTg regardless of diet. (E) At 7 and 12 mo, TNFα was higher in the 3xTg-AD mice than in the NonTg mice in plasma and the Ch-diet increased this TNFα even further. **(F)** Cortical TNFα is increased by the Ch-diet in both genotypes; 3xTg-AD mice had higher TNFα than NonTg mice. **(G-N)** Correlations between plasma choline, cortex choline, ACh, and TNFα in NonTg mice and 3xTg-AD mice at 12 mo. Data are reported as means ± SEM. *P < 0.05, ***P < 0.001, ****P < 0.0001.

We next examined TNFα in plasma and cortex. We found a significant main effect of age, indicating that plasma TNFα increases with age (Fig 2E; *F*_1,20_ = 5.614, *P* = 0.0280). At 7 months, there were significant main effects of both genotype (*F*_1,20_ = 281.0, *P* < 0.0001) and diet (*F*_1,20_ = 9.462, *P* = 0.0060); 3xTg-AD mice had higher levels than NonTg mice and the Ch-diet groups had higher levels than ChN mice. There was also a significant genotype by diet interaction (*F*_1,20_ = 10.80, *P* = 0.0037), where the 3xTg-AD Ch-mice (*P* = 0.0013) had higher plasma TNFα than their ChN counterparts, but there was no significant diet difference between the NonTg mice. In plasma at 12 months of age, we found significant main effects of both genotype (*F*_1,20_ = 4754, *P* < 0.0001) and diet (*F*_1,20_ = 259.7, *P* < 0.0001), where the 3xTg-AD mice and the Ch-mice showed elevated TNFα. We also found a significant genotype by diet interaction (*F*_1,20_ = 168.3, *P* < 0.0001), where 3xTg-AD ChN mice had higher TNFα than both NonTg ChN (*P* < 0.0001) and Ch-(*P* < 0.0001) mice. Further, 3xTg-AD Ch-mice had significantly elevated TNFα compared to their ChN counterparts (*P* < 0.0001). In the cortex (Fig 2F), we also observed significant main effects of both genotype (*F*_1,27_ = 247.2, *P* < 0.0001) and diet (*F*_1,27_ = 64.67, *P* < 0.0001) for TNFα, with 3xTg-AD mice and the Ch-mice exhibiting elevations. Collectively, this data highlights that the 3xTg-AD mice have higher TNFα than NonTg mice at both 7 and 12 months of age, and that the Ch-diet increases levels of this pro-inflammatory cytokine.

We next performed correlation analyses of our plasma and cortical measures in 12-month-old mice. Given the substantial genotypical group differences in ACh and TNFα, our correlation analyses were separated by genotype. We found significant positive correlations between plasma and cortical choline in both the NonTg (Fig. 2G; *r*_10_ = 0.8911, *P* < 0.0001) and 3xTg-AD mice (Fig. 2H; *r*_10_ = 0.8673, *P* = 0.0003). In the NonTg mice, there was a significant positive correlation between plasma choline and ACh (Fig. 2I; *r*_10_ = 0.7854, *P* = 0.0025), but was non-significant in 3xTg-AD mice (Fig. 2J). In both NonTg (Fig. 2K; *r*_10_ = -0.9431, *P* < 0.0001) and 3xTg-AD (Fig. 2L; *r*_10_ = -0.8854, *P* < 0.0001) mice, there were significant negative correlations between plasma choline and TNFα. There was also a significant negative correlation between ACh (Fig. 2M; *r*_10_ = -0.8803, *P* = 0.0002) and TNFα for NonTg mice, but a non-significant correlation in 3xTg-AD mice (Fig 2N). These results highlight the association between choline, its derivate ACh and TNFα, and how 3xTg-AD mice exhibit increased dysregulation of circulating and cortical levels with the Ch-diet.

### Dietary choline deficiency exacerbates pathology in 3xTg-AD mice

Next, we examined the impact of a Ch-diet on pathology in the cortex of 3xTg-AD mice. For Aβ pathology, we assessed soluble and insoluble Aβ (40 and 42), isoforms that aggregate to form Aβ plaques^8, 23^. Our previous report showed that a Ch-diet did not exacerbate soluble Aβ_40_, so this was not measured in the current study^8^. For tau pathology, we analyzed phosphorylated tau epitopes that are pathological in AD, threonine (Thr) 181 and serine (Ser) 396^8, 24, 25^. NonTg mice were excluded because they do not harbor familial mutations leading to AD. We found significant elevations of soluble Aβ_42_ (Fig. 3A; *t*_10_ = 3.134, *P* = 0.0106), insoluble Aβ_40_ (Fig. 3B, left; *t*_10_ = 16.46, *P* < 0.0001) and Aβ_42_ (Fig. 3B, right; *t*_10_ = 16.83, *P* < 0.0001) in 3xTg-AD Ch-mice. We also found significantly higher soluble pTau Thr 181 (Fig. 3C, left; *t*_10_ = 19.12, *P* < 0.0001), insoluble pTau Thr 181 (Fig. 3C, right; *t*_10_ = 18.76, *P* < 0.0001), soluble pTau Ser 396 (Fig. 3D, left; *t*_10_ = 13.58, *P* < 0.0001), and insoluble pTau Ser 396 (Fig. 3D, right; *t*_10_ = 7.053, *P* < .0001) in 3xTg-AD Ch-mice. Collectively, these results highlight that a Ch-diet exacerbates AD pathology, consistent with our previous report^8^.

Next, we performed correlations of the AD pathologies in 3xTg-AD mice. For soluble Aβ_42_, there was no significant correlation with plasma choline (Fig. 3E), but a significant negative correlation with cortical choline (Fig. 3F; *r*_10_ = -0.6561, *P* = 0.0205). For insoluble Aβ_40_, there were significant negative correlations for both plasma (Fig. 3G; *r*_10_ = -0.8398, *P* = 0.0006) and cortical choline (Fig. 3H; *r*_10_ = -0.9825, *P* < 0.0001). For insoluble Aβ_42_, there were significant negative correlations for both plasma (Fig. 3I; *r*_10_ = -0.8329, *P* = 0.0008) and cortical choline (Fig. 3J; *r*_10_ = -0.9706, *P* < 0.0001). For soluble pTau Thr 181, there were significant negative correlations for both plasma (Fig. 3K; *r*_10_ = -0.8766, *P* = 0.0002) and cortical choline (Fig. 3L; *r*_10_ = -0.9810, *P* < 0.0001). For insoluble pTau Thr 181, there were significant negative correlations for both plasma (Fig. 3M; *r*_10_ = -0.8158, *P* = 0.0012) and cortical choline (Fig. 3N; *r*_10_ = -0.9764, *P* < 0.0001). Additionally, for soluble pTau Ser 396, there were significant negative correlations for both plasma (Fig. 3O; *r*_10_ = -0.8813, *P* = 0.0002) and cortical choline (Fig. 3P; *r*_10_ = -0.9708, *P* < 0.0001). Lastly, for insoluble pTau Ser 396, there were significant negative correlations for both plasma (Fig. 3Q; *r*_10_ = -0.7858, *P* = 0.0024) and cortical choline (Fig. 3R; *r*_10_ = -0.9217, *P* < 0.0001). There were no significant correlations between plasma ACh and pathological markers.

**Figure 3:**
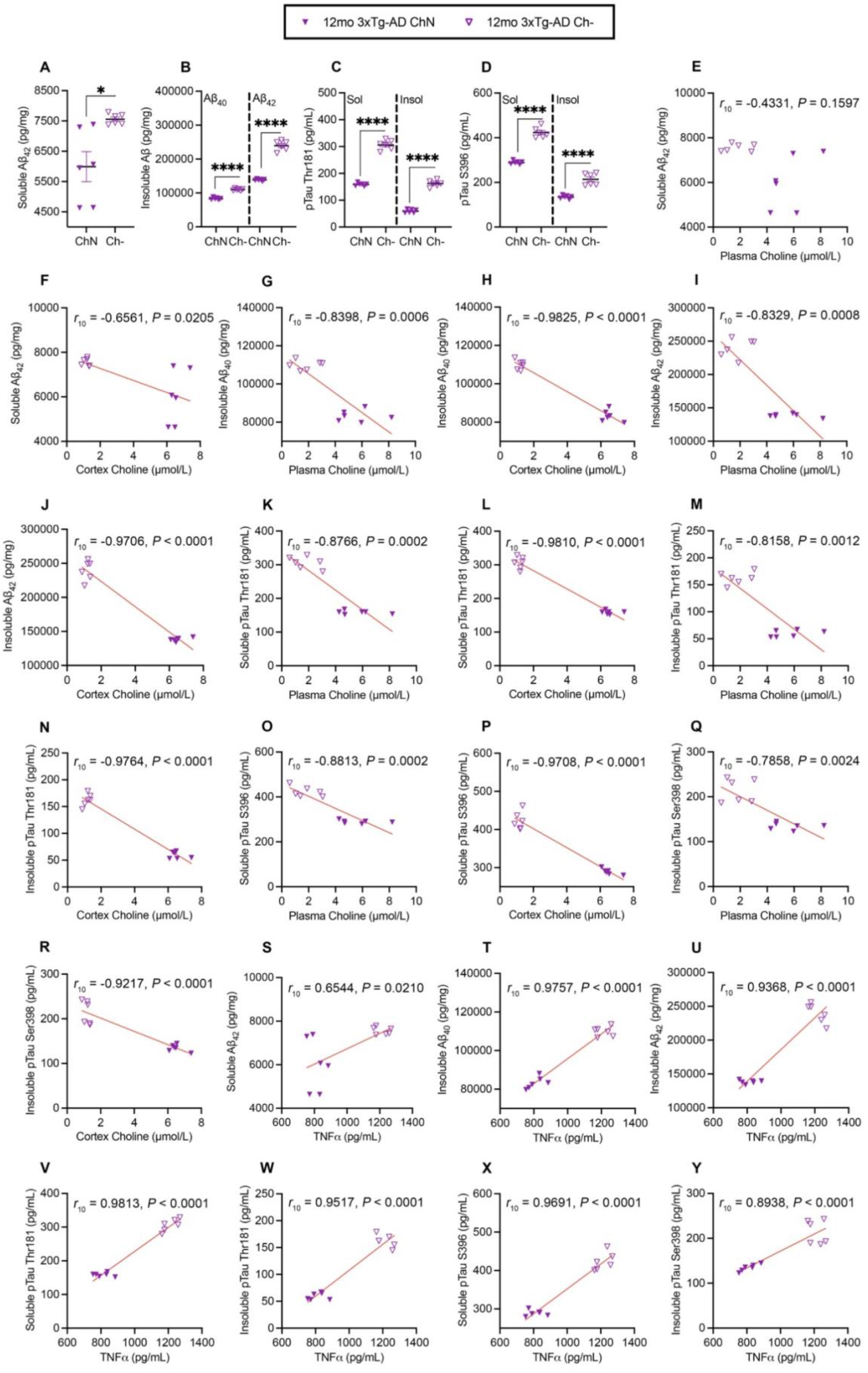
Correlations between choline, ACh, TNFα and the various forms of amyloid and pathological tau epitopes in choline deficient mice. **(A-D)** In the cortex of 3xTg-AD mice, the Ch-diet elevated soluble Aβ_42_, insoluble Aβ_40_, insoluble Aβ_42_, soluble pTau Thr 181, insoluble pTau Thr 181, soluble pTau Ser 396, and insoluble pTau Ser 396. **(E-R)** Correlations between free plasma choline, cortex choline, and soluble Aβ_42_, insoluble Aβ_42_, insoluble Aβ_40_, soluble pTau Thr 181, insoluble pTau Thr 181, soluble pTau Ser 396, and insoluble pTau Ser 396. **(S-Y)** Correlations between plasma TNFα and soluble Aβ_42_, insoluble Aβ_42_, insoluble Aβ_40_, soluble pTau Thr 181, insoluble pTau Thr 181, soluble pTau Ser 396, and insoluble pTau Ser 396.

Conversely, there were significant positive correlations between TNFα and soluble Aβ_42_ (Fig. 3S; *r*_10_ = 0.6544, *P* = 0.0210), insoluble Aβ_40_ (Fig. 3T; *r*_10_ = 0.9757, *P* < 0.0001) and insoluble Aβ_42_ (Fig. 3U; *r*_10_ = 0.9368, *P* < 0.0001). Additionally, there were significant positive correlations between TNFα and both soluble (Fig. 3V; *r*_10_ = 0.9813, *P* < 0.0001) and insoluble (Fig. 3W; *r*_10_ = 0.9517, *P* < 0.0001) pTau Thr 181. Lastly, there was a significant positive correlation between TNFα and both soluble (Fig. 3X; *r*_10_ = 0.9691, *P* < 0.0001) and insoluble pTau Ser 396 (Fig. 3Y; *r*_10_ = 0.8938, *P* < 0.0001). These results highlight that a Ch-diet in the 3xTg-AD mice increased AD pathology that are consistent with changes observed in human AD specimens, reflecting the impact of choline deficiency in disease progression.

### Choline supplementation in the APP/PS1 mouse increases choline and acetylcholine in the brain while reducing TNFα and amyloidosis

To determine the effects of dietary choline supplementation throughout life, APP/PS1 and NonTg mice were placed on a choline supplemented (Ch+) or standard control (Ctl) diet from 2.5 to 12.5 months of age. We then measured brain levels of choline, ACh and TNFα (Fig. 4A). For choline, we found significant main effects of genotype (Fig. 4B; *F*_1,19_ = 39.95, *P* < 0.0001) and diet (*F*_1,19_ = 78.85, *P* < 0.0001), where APP/PS1 mice showed reductions compared to NonTg mice, and Ch+ mice showed elevations compared to their Ctl counterparts. We also found a significant genotype by diet interaction (*F*_1,19_ = 14.45, *P* = 0.0012), where the APP/PS1 Ch+ mice showed higher brain choline than APP/PS1 Ctl (*P* = 0.0066) and NonTg Ch+ show higher levels than NonTg Ctl (*P* < 0.0001). Notably, the brain choline in NonTg Ctl and APP/PS1 Ch+ (*P* = 0.488) did not significantly differ, illustrating a rescue effect. For brain ACh, we found significant main effects of genotype (Fig. 4C; *F*_1,19_ = 42.37, *P* < 0.001) and diet (*F*_1,19_ =108.6, *P* < 0.0001), where APP/PS1 mice showed reductions compared to NonTg mice, and Ch+ mice showed elevations compared to their Ctl counterparts. For brain TNFα, we found significant main effects of genotype (Fig. 4D; *F*_1,19_ = 270.9, *P* < 0.0001) and diet (*F*_1,19_ = 76.75, *P* < 0.0001), where APP/PS1 mice showed elevations compared to NonTg mice, and Ch+ mice showed reductions compared to their Ctl counterparts. We also found a significant genotype by diet interaction (*F*_1,19_ = 120.9, *P* < 0.0001), where the APP/PS1 Ch+ mice showed significantly lower TNFα than APP/PS1 Ctl (*P* < 0.0001). These results illustrate that supplementing with choline can produce protective effects in the brain.

**Figure 4:**
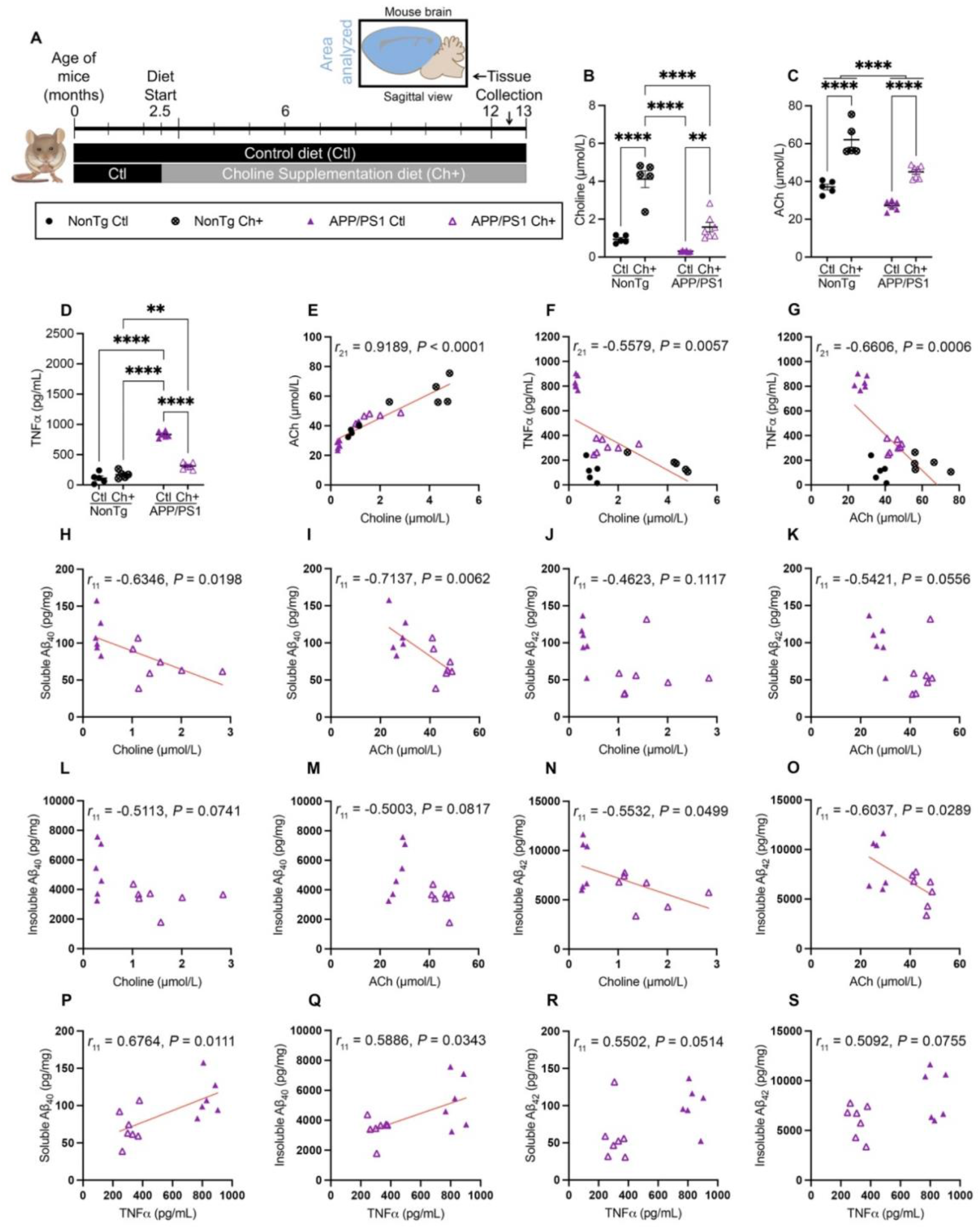
Choline supplementation in the APP/PS1 mouse increases choline and acetylcholine in the brain while reducing TNFα and amyloidosis. **(A)** Timeline of dietary choline manipulation. **(B, C)** APP/PS1 show reduced levels of choline and ACh, and the Ch+ diet increased the levels in both NonTg and APP/PS1 mice. **(D)** APP/PS1 show increased levels of brain TNFα, and the Ch+ diet decrease the levels. **(E-G)** Correlations between choline, ACh and TNFα. **(H-S)** Correlations between brain choline, ACh, TNFα and soluble Aβ_40_ and Aβ_42_, and insoluble Aβ_40_ and Aβ_42._ Data are reported as means ± SEM. **P < 0.01, ***P < 0.001, ****P < 0.0001.

We next performed correlation analyses between our brain measures in APP/PS1 and NonTg mice. We found a significant positive correlation between brain choline and ACh (Fig. 4E; *r*_21_ = 0.9189, *P* < 0.0001). We also found significant negative correlations both between choline (Fig. 4F; *r*_21_ = -0.5579, *P* = 0.0057), ACh (Fig. 4G; *r*_21_ = -0.6606, P = 0.0006), and TNFα. Next, we performed correlations of soluble and insoluble Aβ (40 and 42) in the APP/PS1. For soluble Aβ_40_, there were significant negative correlations with both choline (Fig. 4H; *r*_11_ = -0.6346, *P* = 0.0198), and ACh (Fig. 4I; *r*_11_ = -0.7137, *P* = 0.0062). There was no significant correlation between soluble Aβ_42_ and choline (Fig. 4J), but a trend toward significance in the correlation between soluble Aβ_42_ and ACh (Fig. 4K; *r*_11_ = -0.5421, *P* = 0.0556). Additionally, there were no significant correlations between insoluble Aβ_40_ and both choline and ACh (Fig. 4L, M). For insoluble Aβ_42,_ there were significant negative correlations with both choline (Fig. 4N; *r*_11_ = -0.5532, *P* = 0.0499), and ACh (Fig. 4O; *r*_11_ = -0.6037, *P* = 0.0289). Lastly, there were significant positive correlations between TNFα and soluble Aβ_40_ (Fig. 4P; *r*_11_ = 0.6764, *P* = 0.0111) and insoluble Aβ_40_ (Fig. 4Q; *r*_11_ = 0.5886, *P* = 0.0343). There was a trending positive correlation between TNFα and soluble Aβ_42_ (Fig. 4R), but no correlation with insoluble Aβ_42_ (Fig. 4S). These results highlight the association between choline, ACh, and TNFα, and how a Ch+ diet in APP/PS1 can reduce the levels of various forms of soluble and insoluble amyloid fractions.

### Metabolomics analysis reveals four key metabolites significantly correlated with choline

To investigate the metabolic differences in human subjects with the varying severities of AD and CONs, we conducted a comprehensive targeted metabolomic profile of metabolites from human serum and probed for associations with choline (Supplemental Fig. 2). Correlation analysis identified four metabolites that were significantly correlated with choline levels (Fig. 5A). There was a significant negative correlation of choline with L-valine (Fig. 5B; *r*_33_ = -0.442, *P* = 0.0179). Three metabolites showed a significant positive correlation with choline, including 4-hydroxyphenylpyruvic acid (4-HPPA; Fig 5C; *r*_34_ = 0.458, *P* = 0.0056), Methylmalonic acid (MMA; Fig 5D; *r*_34_ = 0.3653, *P* = 0.0285), and Ferulic acid (Fig 5E; *r*_34_ = 0.3333, *P* = 0.047). We next examined the correlation of ACh with the metabolites and found a significant negative correlation of ACh with L-valine (Fig. 5F; *r*_33_ = -0.488, *P* = 0.0029) and positive correlations of ACh with 4-HPPA (Fig. 5G; *r*_34_ = 0.4066, *P* = 0.0139). The correlation between ACh and MMA was non-significant (Fig. 5H). There was a significant positive correlation between ACh and Ferulic acid (Fig. 5I; *r*_34_ = 0.3806, *P* = 0.022). Collectively, these results provide novel insights into the metabolic alterations associated with both choline and ACh levels.

**Figure 5:**
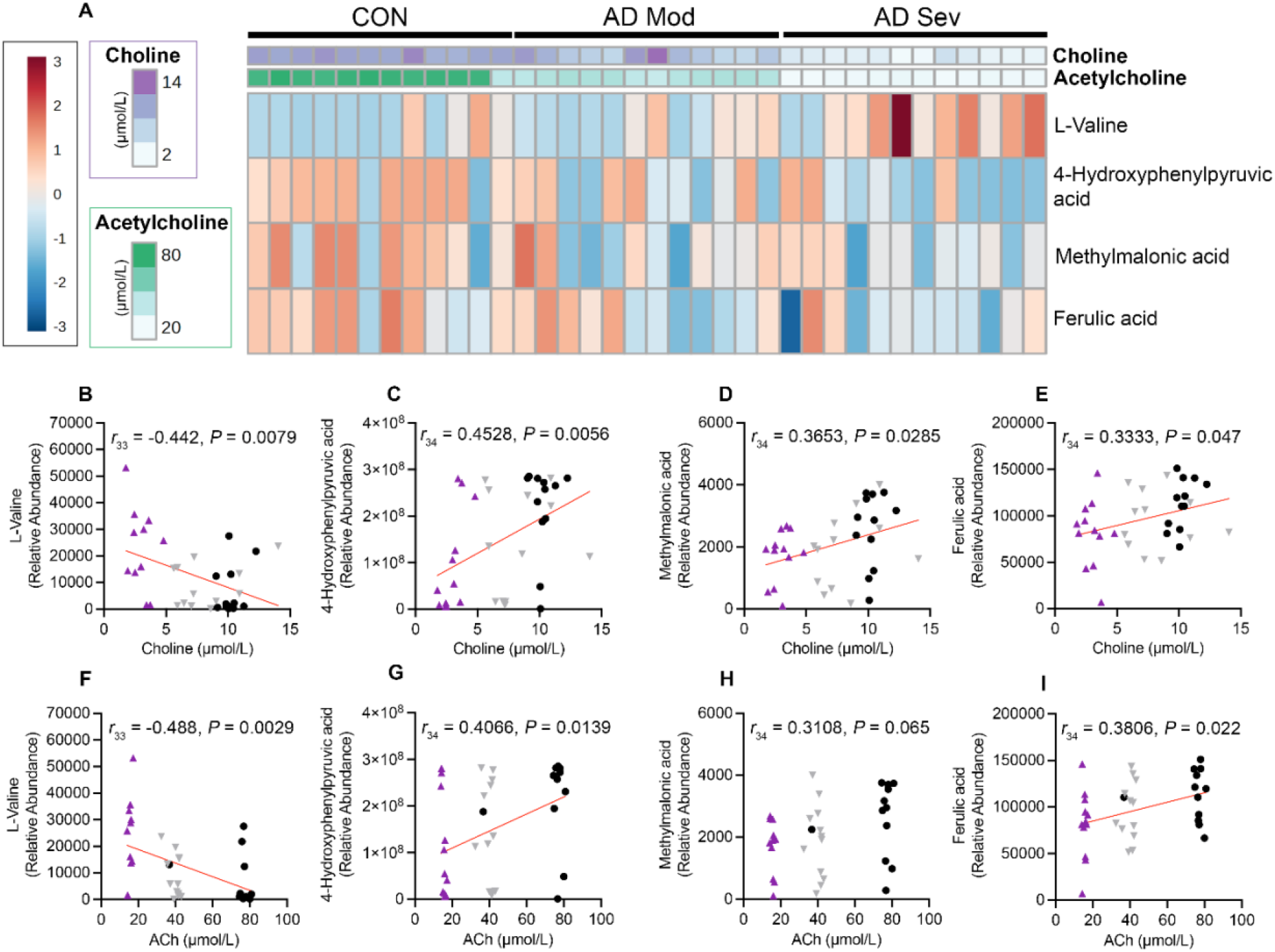
Metabolomic analysis from human serum reveals metabolites that correlate with choline and acetylcholine (ACh). **(A)** Heatmap illustrating the relative abundance of each of the four metabolites significantly correlated with choline per sample. **(B)** A significant negative correlation between circulating choline levels and L-Valine. **(C-E)** Significant positive correlations between circulating choline levels and 4-Hydroxyphenylpyruvic acid, Methylmalonic acid and Ferulic acid. **(F)** A significant negative correlation between circulating ACh levels and L-Valine. **(G-I)** Significant positive correlations between circulating ACh levels and 4-Hydroxyphenylpyruvic acid and Ferulic acid, and a non-significant trend with Methylmalonic acid.

## Discussion

We found disease-associated reductions in serum choline and ACh in AD patients compared to healthy controls. Conversely, serum TNFα levels were elevated in all AD cases compared to healthy controls and the highest levels were associated with highest pathological burden. Additionally, we assessed the relationship between choline and other metrics of brain health. CAA cases displayed lower serum choline and ACh, and higher TNFα. Further, CWMR cases had lower choline than cases without CWMR, illustrating the importance of choline for white matter integrity. Previous research that found relationships between decreased choline intake and increased risk for cognitive decline and AD development estimated intake levels and did not measure pathology ^11,12^Consequently, this is the first study examining the relationship between circulating choline levels and unique disease states, highlighting the importance of monitoring choline for brain health.

We examined the impact of dietary choline deficiency and supplementation on pathogenesis in two mouse models of AD and found parallels to our human findings. Higher TNFα, Aβ, and tau phosphorylation were correlated with lower plasma choline levels due to the dietary deficiency in the 3xTg-AD mice. Notably, the Ch-diet resulted in lower plasma choline and ACh, and elevated TNFα at 7 months, an age when amyloidosis commences in 3xTg-AD mice, but prior to NFT pathology. Thus, dietary choline differences predate and contribute to pathology; at 7 months, there was less plasma choline in the 3xTg-AD mice than the NonTg mice. Conversely, dietary choline supplementation in APP/PS1 mice elevated choline and ACh while reducing both TNFα and Aβ pathological burden in the brain. Together, this data provides evidence that higher levels of circulating choline correspond with reduced AD pathology.

Elevated L-valine, an essential amino acid, has been linked to a variety of disease-associated mechanisms, including oxidative stress and insulin resistance^26^. While elevated L-valine increases the risk of AD in some studies^27^, other studies have shown lower L-valine in AD^28^. Interestingly, a high choline diet reduces circulating L-valine levels, likely through increased catabolism or uptake by muscles^29^. We found that higher choline and ACh are associated with lower L-valine; the lowest L-valine levels were in the CON and AD Mod groups, supporting evidence that reduced L-valine is protective against AD^28^, collectively indicating that reducing L-valine may be a protective mechanism of choline against AD.

Two identified metabolites correlating with choline and ACh levels have anti-acetylcholinesterase activity, which prevent enzymatic breakdown of ACh. 4-HPPA, an amino acid metabolite, has high specificity and inhibitory activity for acetylcholinesterase ^30, 31^. Ferulic acid is a plant-derived antioxidant that’s been shown to improve AD pathology^32, 33^. Ferulic acid also has anti-acetylcholinesterase activity and increases ACh^32, 33^. Interestingly, many existing drugs to treat AD are acetylcholinesterase inhibitors^31^, suggesting these metabolites can be particularly beneficial against AD reductions in ACh. Notably, serum ACh was positively correlated with 4-HPPA and ferulic acid, supporting the anti-acetylcholinesterase activity. Indeed, this data suggests elevated choline may confer dual protective mechanisms; both increasing the available choline for synthesizing ACh and increasing metabolites with anti-acetylcholinesterase activity.

We found a significant positive correlation between MMA and choline levels, but not ACh levels. MMA is a precursor to succinyl-CoA, a key participant in the citric acid cycle, a major source of ATP and energy^34^. The citric acid cycle is dysregulated in AD^35^. We found the lowest choline levels corresponded to lower MMA. Metabolomic studies have found that MMA is decreased in patients with MCI and AD compared to healthy controls^36^. This supports our findings because the AD Sev group had the lowest choline levels. Other evidence suggests that elevated MMA may be a risk factor for disease states^37^, including dementia^38^. While studies show that choline and MMA levels are inversely related^37, 39^, a study on maternal choline supplementation in mice showed a small, but significant increase in MMA in choline supplemented dams^39^, consistent with our findings. While our results are consistent with some results about MMA and its relationship with AD and choline, the literature is mixed, and future studies should further clarify the relationship between MMA and choline in AD.

Other recent studies support our findings that choline is decreased in the serum and brain of AD patients compared to healthy controls ^40–42, 45^. However, others have observed increases in circulating choline in recently diagnosed AD patients^16, 43^, which may reflect neuronal membrane degradation, releasing choline from choline-containing phospholipids (CCPLs)^43^. Reductions in free choline reduces the capacity to maintain both CCPLs and ACh ^44, 45^. Cells prioritize ACh synthesis over CCPLs, leading to “autocannibalism” of CCPLs to free choline for ACh production^44, 46^. Only cholinergic neurons use their CCPLs as choline reserves for ACh production^45^, and it is likely that reserves are diminished with disease progression leading to choline levels reflecting dietary intake and endogenous production. Regardless, increased choline intake may prevent or delay the “autocannibalism” of CCPLs, preserving neuronal integrity.

Our human subject profiles did not include information about dietary habits, which has been documented in previous studies linking low choline with AD^11, 12, 40^. While we didn’t find significant differences in BMI between our human groups, appetite loss is common in AD^47^. Often caregivers of patients choose enjoyable foods over nutritional foods^47^, suggesting that overall calories may be similar between groups, but nutritional values, including choline, might be lower in AD than healthy controls. Notably, an overall reduction of food consumption has been shown to result in reduced circulating choline within days^4^. Reduced circulating choline was clearly associated with greater pathology in the current study, which may be influenced by food consumption, PEMT variation and choline need throughout various organ systems. Future work in humans should establish how dietary choline intake and PEMT variation alters circulating choline levels, allowing for a complete assessment of its contributions to pathogenesis.

In conclusion, low circulating choline levels were associated with increased inflammation and neuropathology, suggesting that both consuming adequate dietary choline and monitoring circulating choline levels should be implemented for healthy aging. While adequate or even supplementation of dietary choline intake may not be able to reduce plaques or tangles when provided at an advanced stage of AD, it is critical for proper brain function throughout adulthood, serving as a preventive strategy to dampen AD pathological progression.

## Materials and Methods

### Human serum samples

Human serum samples were obtained from the Arizona Study of Aging and Neurodegenerative Disorders/Brain and Body Donation Program, as previously described^18, 19^. Thirty-six samples, balanced for sex, were obtained including healthy controls with Braak stage ≤ III (CON, n = 12), moderate AD with Braak stage = IV and moderate to frequent cortical neuritic plaques (AD Mod, n = 12), and severe AD with Braak stage = VI and frequent cortical neuritic plaques (AD Sev, n = 12). AD Mod and AD Sev corresponded with NIA-RI Intermediate and High classifications, respectively^19^. The medical profiles of the subjects were extensively documented both pre- and post-mortem. Profiles included body mass index (BMI), diagnosis age, years since diagnosis, expired age, post-mortem interval (PMI) at autopsy, final Mini-Mental State Examination (MMSE), APOE status, Braak stage, CERAD neuritic plaque density, TDP-43 pathology, brain weight, and National Institute on Aging-Regan Institute (NIA-RI) diagnosis. Missing metrics for some cases are denoted in Supplementary Table 1. Some subjects had TDP-43 proteinopathy but did not meet diagnosis criteria for frontotemporal lobar degeneration with TDP-43. No other neuropathologies were present.

### Animals

Mice were kept on a 12-h light/dark cycle at 23°C with *ad libitum* access to food and water and group-housed, four to five per cage. All animal procedures were approved in advance by the Institutional Animal Care and Use Committee of Arizona State University. 3xTg-AD mice express a Swedish mutation of the amyloid precursor protein (APP), a mutated presenilin (PS1 M146V) to accelerate amyloid deposition, and a mutated human tau (P301L), leading to tau pathology, and were generated as previously described^8, 19, 48^. C57BL6/129Svj mice were used as NonTg controls. Consistent with previous literature highlighting inconsistent pathology in male 3xTg-AD, only female mice were used^8, 49^. 3xTg-AD mice develop Aβ starting at 6 months and pathological tau at 12 months^8, 19, 49^. 3xTg-AD and NonTg mice were randomly assigned to one of two diets at 3 months of age; a standard laboratory AIN76A diet (Envigo Teklab Diets, Madison WI) with adequate choline (ChN; 2.0 g/kg choline chloride; #TD.180228), or a AIN76A choline-deficient (Ch-; 0.0 g/kg; #TD.110617) diet, as previously described^8^.

APP/PS1 mice were generated for the choline supplementation experiment as previously described^15^. The APP/PS1 mice are hemizygous for the amyloid precursor protein (APP) Swedish mutations (KM670/671Nl) and presenilin1 (PS1) deltaE9 mutation. APP/PS1 mice were backcrossed for 12 generations into a pure 129/SvJ background^15^. APP/PS1 mice develop Aβ_42_ plaques starting at 6 months, with extensive pathology through the hippocampus and cortex at 12 months^15, 50, 51^. At 2.5 months of age, female APP/PS1 and wild-type (NonTg) mice were randomly assigned to receive one of two concentrations of choline chloride in their diet (Harlan laboratories). The control diet (Ctl; 1.1 g/kg choline chloride) supplies adequate choline and is comparable to amounts in previous studies^52–54^, while the supplemented diet (Ch+; 5.0 g/kg choline chloride) contains approximately 4.5 times the amount of choline consumed in the Ctl group, allowing assessment of the effects of supplementation.

### Mouse tissue and plasma collection

At 7 and 12 months of age, 3xTg-AD and NonTg mice were fasted for 16 hours, and 150– 200 μl (≤ 10% of the subject’s body weight) of blood was collected via the submandibular vein and placed into EDTA-lined tubes (BD K_2_EDTA #365974). Tubes were inverted eight times to assure anticoagulation, kept on ice for 60–90 min, and then centrifuged at 2200 RPM for 30 min at 4°C to separate phases. The top layer was collected and frozen at −80°C prior to use. Plasma samples were used to assess choline, acetylcholine (ACh), and tumorous necrosis factor α (TNFα). 3xTg-AD and NonTg mice were euthanized at a mean age of 12.7 months via perfusion with 1x PBS or CO2 inhalation. Cortex tissue was dissected, homogenized to extract soluble and insoluble fractions, and stored at -80°C, as previously described^8^. APP/PS1 and NonTg counterparts were euthanized at 12.5 months of age via perfusion with 1x PBS or CO2 inhalation, olfactory bulbs and cerebellum were removed, and the remaining brain hemisphere was separated, homogenized to extract soluble and insoluble fractions, and stored at –80°C, as previously described^15^.

### ELISAs and colorimetric assays

We used commercially available ELISA kits (Invitrogen-ThermoFisher Scientific) to quantify levels of soluble and insoluble Aβ_42_ and insoluble Aβ_40_, and levels of phosphorylated Tau (pTau) at threonine (Thr) 181 and serine (Ser) 396, and TNFα (Abcam, ab208348) as previously described^8, 55^. Choline and acetylcholine (ACh) levels in serum, plasma and cortex were quantified using commercially available colorimetric assay kits (Abcam, ab219944; Cell Biolabs, Inc., STA-603). The ACh assay raw value is a measure of total choline, both the choline bound in ACh and free choline. Consequently, the value for ACh that was analyzed was derived from the formula Aceytalcholine = Total Choline − Free Choline, as described by the manufacturer.

### Metabolomics analysis

Analytical methanol (MeOH) was purchased from Fisher Scientific (Waltham, MA). Methyl tert-butyl ether (MTBE), *O*-methyl hydroxylamine hydrochloride (MeOX), N-Methyl-N-(tert-butyldimethylsilyl) trifluoroacetamide (MTBSTFA), and pyridine were purchased from Sigma-Aldrich (Saint Louis, MO). Deionized water was provided in-house by a water purification system from EMD Millipore (Billerica, MA). Phosphate buffered saline (PBS) was purchased from GE Healthcare Life Sciences (Logan, UT). Compound standards were purchased from Sigma-Aldrich and Avanti Polar Lipids (Alabaster, AL).

Experimental samples were prepared for GC-MS analysis as previously described^56^. Human serum samples were thawed at 4°C. Next, 200 μL 10x diluted PBS and 80 μL of MeOH containing 50 μM PC (17:0, 17:0) and PG (17:0, 17:0) internal standards were added to each thawed sample (20 μL for human serum). Afterward, 400 μL of MTBE was added to each sample (MTBE:MeOH:H_2_O = 10:2:5, v/v/v), vortexed for 30 sec, and then stored at –20°C for 20 min. Lastly, samples were centrifuged at 21,300 *g* to separate aqueous and MTBE phases. The aqueous bottom layer (180 μL) from the MTBE extraction described above was collected into new Eppendorf tubes for derivatization prior to targeted metabolic profiling with GC-MS. The aqueous layer was dried under vacuum at 37°C for 4 h using a Savant SpeedVac vacuum concentrator. The residues were first derivatized with 40 µL of 20 mg/mL MeOX solution in pyridine under 60°C for 90 min. Next, 60 µL of MTBSTFA containing d_27_-mysristic acid were added, and the mixture was incubated at 60°C for 30 min. The samples were then vortexed for 30 sec, followed by centrifugation at 21,300 *g* for 10 min. Finally, 70 µL of supernatant were collected from each sample into new glass vials for GC-MS analysis while 10 µL of each sample was pooled to create a quality control (QC) sample, which was injected once every 10 experimental samples to monitor gradual changes in systems performance.

GC-MS conditions used here were mainly adopted from previous studies^57, 58^. Briefly, GC-MS experiments were performed on an Agilent 7820A GC-5977B MSD system (Santa Clara, CA) by injecting 1 µL of prepared samples. Research-grade helium (99.9995% purity) was used as the carrier gas with a constant flow rate of 1.2 mL/min. Front inlet, auxiliary line, and source temperatures were set to 250°C, 290°C, and 230°C, respectively. The separation of metabolites was achieved using an Agilent HP-5ms capillary column (30 m x 250 µm x 0.25 µm). The column temperature was maintained at 60°C for 1 min, increased at a rate of 10°C/min to 325°C, and then held at this temperature for 10 min. Data were recorded following a 3 min solvent delay. Electron energy was set to –70 eV, and mass spectral data were collected between *m/z* 60-550. Data extraction was performed using Agilent MassHunter Profinder software. Collected data were unbiasedly queried against an internal spectral and retention time library for the identification of the 126 targeted analytes and an RT tolerance of 0.10 min was used.

### Statistical analysis

Data was analyzed using GraphPad Prism version 9.5.1 (GraphPad Software). Group differences for human serum samples were analyzed using a one-way analysis of variance (ANOVA) followed by Bonferroni’s corrected post hoc test, when appropriate. Unpaired t-tests were used to analyze differences between the human AD groups and between the diet groups, when appropriate. Group differences for mouse samples were analyzed using two-way factorial ANOVAs (for genotype by diet). Linear correlations were calculated using the Pearson’s r coefficient. For metabolomics analysis, following peak integration, metabolites were filtered for reliability and only those with QC coefficient of variation (CV) < 20% and relative abundance of 1,000 in > 80% of samples were retained for statistical analysis. Univariate and multivariate analyses of metabolomics data were performed using the MetaboAnalystR package^57^. A total of 184 missing values (4.7%) were detected in the human serum dataset. Metabolites with >50% missing values were removed from analysis, and remaining missing values were imputed using a sample-wise k-nearest neighbors’ algorithm. Human serum samples were sum normalized, were log_10_-transformed, and mean centered before analysis. Significance was set at *P* < 0.05.

## Acknowledgments

We would like to thank Savannah Tallino for assisting with figure design. We are grateful to the Banner Sun Health Research Institute Brain and Body Donation Program of Sun City, Arizona for the provision of human biological materials.

## Author contributions

Conceptualization: RV, JMJ

Investigation: JMJ, WW, PJ

Visualization: JMJ, RV, PJ

Supervision: RV, JKS

Writing—original draft: JMJ

Writing—review & editing: JMJ, PJ, WW, GES, TGB, JKS, RV

## Data and materials availability

Data are available in the main text or the supplementary materials. Raw data supporting the conclusions of this work will be made available upon reasonable request.

## Supplementary Materials

**Supplemental Figure 1:**
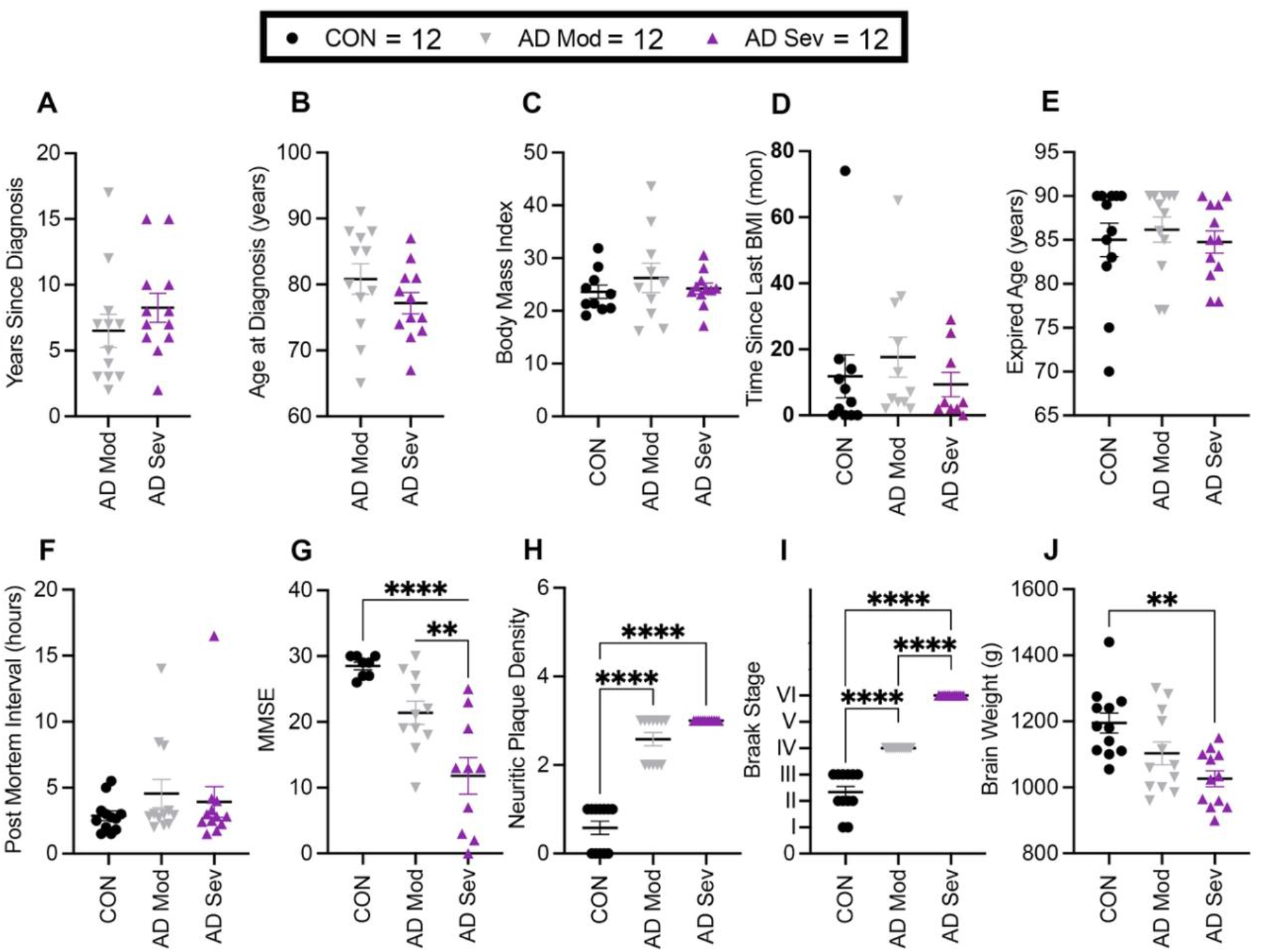
Graphical representation of human subject profiles. Subjects were matched on **(A)** years since diagnosis, **(B)** age at diagnosis, **(C)** BMI, **(D)** time since last BMI measurement, **(E)** expired age, and **(F)** PMI. Subjects differed from each other according to diagnosis on **(G)** MMSE, **(H)** plaque density, **(I)** Braak stage, **(J)** brain weight. Data are reported as means ± SEM. *p<0.05, **p<0.01, ****p<0.0001.

**Supplementary Table 1: Human characteristic profiles.**

**Supplemental Figure 2: Raw data from the metabolomic analysis.**

